# Structural basis for anomalous cellular trafficking behavior of glaucoma-associated A427T mutant myocilin

**DOI:** 10.1101/2025.02.26.640437

**Authors:** Kamisha R. Hill, Hailee F. Scelsi, Hannah A. Youngblood, Jennifer A. Faralli, Tatsuo Itakura, M. Elizabeth Fini, Donna M. Peters, Raquel L. Lieberman

## Abstract

Familial mutations in myocilin cause vision loss in glaucoma due to misfolding and a toxic gain of function in a senescent cell type in the anterior eye. Here we characterize the cellular behavior and structure of the myocilin (myocilin ^A427T^) mutant, of uncertain pathogenicity. Our characterization of A427T demonstrates that even mutations that minimally perturb myocilin structure and stability can present challenges for protein quality control clearance pathways. Namely, when expressed in an inducible immortalized trabecular meshwork cell line, inhibition of the proteasome reroutes wild-type myocilin, but not myocilin ^A427T^, from endoplasmic reticulum associated degradation to lysosomal degradation. Yet, the crystal structure of the A427T myocilin olfactomedin domain shows modest perturbations largely confined to the mutation site. The previously unappreciated range of mutant myocilin behavior correlating with variable stability and structure provides a rationale for why it is challenging to predict causal pathogenicity of a given myocilin mutation, even in the presence of clinical data for members of an affected family. Comprehending the continuum of mutant myocilin behavior in the laboratory supports emerging efforts to use genetics to assess glaucoma risk in the clinic. In addition, the study supports a therapeutic strategy aimed at enhancing autophagic clearance of mutant myocilin.

**Significance statement:**

- Rare familial mutations cause early onset glaucoma
- A427T is a case of uncertain pathogenicity
- A427T is structurally similar to wild-type but is not efficiently degraded

## Introduction

Approximately one third of the human proteome is folded in the endoplasmic reticulum (ER). An extensive protein homeostasis network ensures that proteins in the ER are synthesized, properly folded, and trafficked to their final destinations. If folding checks fail, proteins are targeted for efficient degradation by ER-associated degradation (ERAD) by unfolded protein response (UPR) signaling pathways (Klaips *et al*., 2018; Karagoz *et al*., 2019; Kettel and Karagoz, 2024). In this way, the ER can respond to errors during synthesis and folding to minimize protein aggregation and maintain cellular protein levels. The efficiency of the proteostasis machinery declines with age, however, resulting in accumulation of protein aggregates that leads to disease (Hipp *et al*., 2019). Proteostasis decline and disease onset are accelerated in cases of mutations in disease-associated genes (Wang and Moult, 2001; Bershtein *et al*., 2013; Santra *et al*., 2019).

Glaucoma, the leading cause of irreversible blindness worldwide, is a heterogeneous collection of neurodegenerative ocular diseases that affect more than 70 million individuals worldwide (Tham *et al*., 2014; Allison *et al*., 2020). The primary risk factor, elevated intraocular pressure (IOP), is brought about by trabecular meshwork (TM) cell and tissue dysfunction in the anterior eye. Similar to neurons, TM cells are long lived and senescent (Liton *et al*., 2005) and as such are likely similarly programmed to avoid programmed cell death (Wang, 1995).

Missense mutations in the gene encoding for myocilin (*MYOC*, Uniprot accession **<u>Q99972</u>**) are associated with a Mendelian-inherited subtype of glaucoma, accounting for 3-4% of primary open angle glaucoma (POAG) and 10-30% of juvenile open angle glaucoma (JOAG) (Stone *et al*., 1997; Fingert *et al*., 1999; Resch and Fautsch, 2009). Wild-type (WT) myocilin (myocilin^WT^) is secreted from TM cells whereas disease-associated myocilin mutants, instead of being degraded, aggregate in the ER, causing a cytotoxic gain of function (Jacobson *et al*., 2001; Joe *et al*., 2003; Liu and Vollrath, 2004; Gould *et al*., 2006; Yam *et al*., 2007).

Myocilin-associated glaucoma falls under the umbrella of proteostasis diseases. Myocilin is a large multidomain protein, yet >90% of the known mutations are clustered within its C-terminal 30 kDa olfactomedin (OLF) domain spanning amino acids 245–504 (Donegan *et al*., 2015). There are many different rare pathogenic mutations, found in populations around the world (Sun *et al*., 2022). In the clinic, pathogenic mutations are characterized by a multigenerational autosomal dominant inheritance pattern with high penetrance, high IOP, specific optic nerve deformities, and age at diagnosis < 40 years (Gordon *et al*., 2002). In the lab, pathogenic mutations exhibit decreased protein stability, increased aggregation in the ER, and multiple hallmarks of amyloid (Burns *et al*., 2010; Burns *et al*., 2011; Orwig *et al*., 2012; Hill *et al*., 2014; Lieberman and Ma, 2021). Not all myocilin mutations are pathogenic; benign polymorphisms, usually identified in population-based genomic screening, are often indistinguishable from wild-type myocilin in structure, stability, and cellular behavior (Scelsi *et al*., 2023). These mutations, however, are generally rare, often with allele frequency of ∼1×10^−6^ or corresponding allele counts of ∼10 in the gnomAD database (Chen *et al*., 2024).

Here, we focus on a myocilin variant of uncertain pathogenicity, myocilin^A427T^. We show that despite generally harboring several near-native features, myocilin^A427T^ is at a tipping point for quality control machinery fidelity. Placed in the context of clinical and laboratory literature on myocilin mutants, our study demonstrates that cellular behavior can be affected for variants even with minor biophysical perturbations. This work helps explain why it can be challenging to predict disease causality of a given myocilin mutation even in the presence of clinical data for members of an affected family. Our results also support emerging efforts to use genetics to assess glaucoma risk in the clinic and should prompt new efforts to enhance autophagic clearance of mutant myocilin as a therapeutic strategy.

## Results & Discussion

### Pathogenicity of A427T myocilin variant is uncertain

The myocilin missense variant A427T, was first identified in the French-Canadian population in Quebec (Faucher *et al*., 2002). Initially published as a sporadic case of POAG diagnosed at 73, the mutation was later found to be a familial case with autosomal dominant inheritance. Other family members that carried the A427T mutation around the same age as the proband were found to be symptomatic but diagnosed with other types of glaucoma, and several younger family members were asymptomatic despite carrying the mutation. More sporadic cases of POAG linked to A427T were identified in South Asian individuals through genetic screens, with no detailed family studies (Bhattacharjee *et al*., 2007; Banerjee *et al*., 2012). Of the two sporadic case individuals, one was diagnosed at 76 with POAG and Parkinson’s disease and the other was diagnosed with POAG at 46 (Bhattacharjee *et al*., 2007). The gnomAD database (https://gnomad.broadinstitute.org/), which does not include clinical data, lists A427T with a total allele frequency of 2.23e-5, and most prevalent in South Asian populations (allele frequency 1.43e-4) (**Table 1**) (Chen *et al*., 2024). Overall, A427T is a rare mutation and could be pathogenic.

**Table 1.** Clinical and population-based data of myocilin^A427T^.

| ClinVar and gnomAD Data | Population | Allele frequency (count) | Clinical Sources | (number of) Individual diagnosis and age at diagnosis |
| --- | --- | --- | --- | --- |
| Accession: **VCV001686783.3**<br>Classification: **Uncertain Significance**<br>Frequency: **0.00002230 (36)** | European (non-Finnish) | 0.00001610 (19) | Faucher, 2002 | (1) POAG; (1) NTG; (1) ACG; (3) asymptomatic*<br>73; 68; 77; (40.5) |
|  | South Asian | 0.0001427 (13) | Bhattacharjee, 2007<br>Banerjee, 2012 | (1) POAG, PD (1) POAG<br>76; 46 |
|  | Admixed American | 0.00003334 (2) | n/a | n/a |
|  | Other (remaining) | 0.00003201 (2) | n/a | n/a |

Data referenced from gnomAD v4.1.0 and ClinVar's September 30, 2024 release.
\*POAG, primary open-angle glaucoma; NTG, normal tension glaucoma; ACG, angle closure glaucoma; PD, Parkinson's Disease; (), median age.

When the isolated OLF domain of myocilin^A427T^ (OLF^A427T^) is recombinantly expressed and purified, it exhibits perplexing biophysical properties distinct from both wild type OLF (OLF^WT^) and disease-causing OLF variants (Donegan *et al*., 2012; Hill *et al*., 2014). The thermal stability of OLF^A427T^ is 47.8 °C, just above a statistically robust 47 °C cutoff between wild-type-like OLF variants and pathogenic OLF variants (Scelsi *et al*., 2023). However, when incubated at 36 °C, purified OLF^A427T^ forms Thioflavin T (ThT)-positive (i.e., amyloid-like) aggregates at a rate in between that of OLF^WT^ and *bona fide* pathogenic variant OLF^D380A^ (Hill *et al*., 2014). In line with these data, the ClinVar consortium, which takes an approach combining clinical metrics and to a lesser extent biophysical characteristics (Landrum *et al*.), considers A427T (https://www.ncbi.nlm.nih.gov/clinvar/variation/1686783/) to be a variant of uncertain significance.

### Myocilin ^A427T^ exhibits a mild misfolding phenotype compared to myocilin^WT^

Cellular secretion assays conducted in the laboratory determine the likelihood of pathogenicity of a given myocilin variant by comparing the extent of myocilin in spent media and the relative abundance of intracellular detergent-insoluble (i.e., aggregated) and detergent-soluble (i.e., not aggregated) species, in Western blot (Zhou and Vollrath). In previous studies, we and others have shown that pathogenic myocilin mutants such as P370L, predominantly localize to the detergent-insoluble fraction (Liu and Vollrath; Stothert *et al*.; Scelsi *et al*.). Less severe variants secrete to a lower extent than WT-like but may accumulate more in the soluble than the insoluble fraction (Scelsi *et al*., 2023). Regarding myocilin^A427T^ in particular, overexpression in immortalized human trabecular meshwork (TM-1) (Filla *et al*., 2002) or COS-7 cells has shown myocilin ^A427T^ present in secreted and soluble fractions (Gobeil *et al*., 2006).

To avoid potential pitfalls due to robust overexpression models in cell lines, here we used TM-1 cells that stably express doxycycline (dox) inducible constructs of full length myocilin^WT^ or myocilin^A427T^ to evaluate variant myocilin behavior. These cells were constructed as previously reported and were shown to secrete myocilin^A427T^ (Itakura *et al*., 2015) but were generated anew (see **Materials and Methods**). To test the effect of myocilin levels on secretion, we treated cells with increasing levels of dox (dox, 1, 2, and 5 µg/mL). In all cases, myocilin^A427T^ was secreted to a lesser extent than myocilin^WT^ (**Figure 1A**). In the detergent-soluble fraction, myocilin^A427T^ was present at markedly higher levels than myocilin^WT^ across dox dosages. In the insoluble fraction, myocilin^A427T^ could be detected, albeit to a limited extent, whereas myocilin^WT^ was absent from the insoluble fraction across all concentrations tested. Interestingly, dose-dependent expression levels appeared to have little if any effect on secretion and solubility of either myocilin^WT^ or myocilin^A427T^. Findings from immunoblot are confirmed by immunocytochemistry, which showed a relative increase in total intracellular expression of myocilin^A427T^ compared to myocilin^WT^ (**Figure 1B-C**). Although the increase was not statistically significant, distinctions in subcellular localization are apparent. Compared to the consistently diffuse intracellular expression of myocilin^WT^, myocilin^A427T^ was characterized by both diffuse intracellular expression as well as some dense perinuclear bands, of which the latter may reflect aggregated myocilin^A427T^ seen in Western blot. Overall, the high level of intracellular retention combined with a modest presence in the insoluble fraction are consistent with a mild cellular misfolding phenotype for myocilin^A427T^ and indicate challenges to mutant myocilin turnover in TM cells.

**Figure 1.**
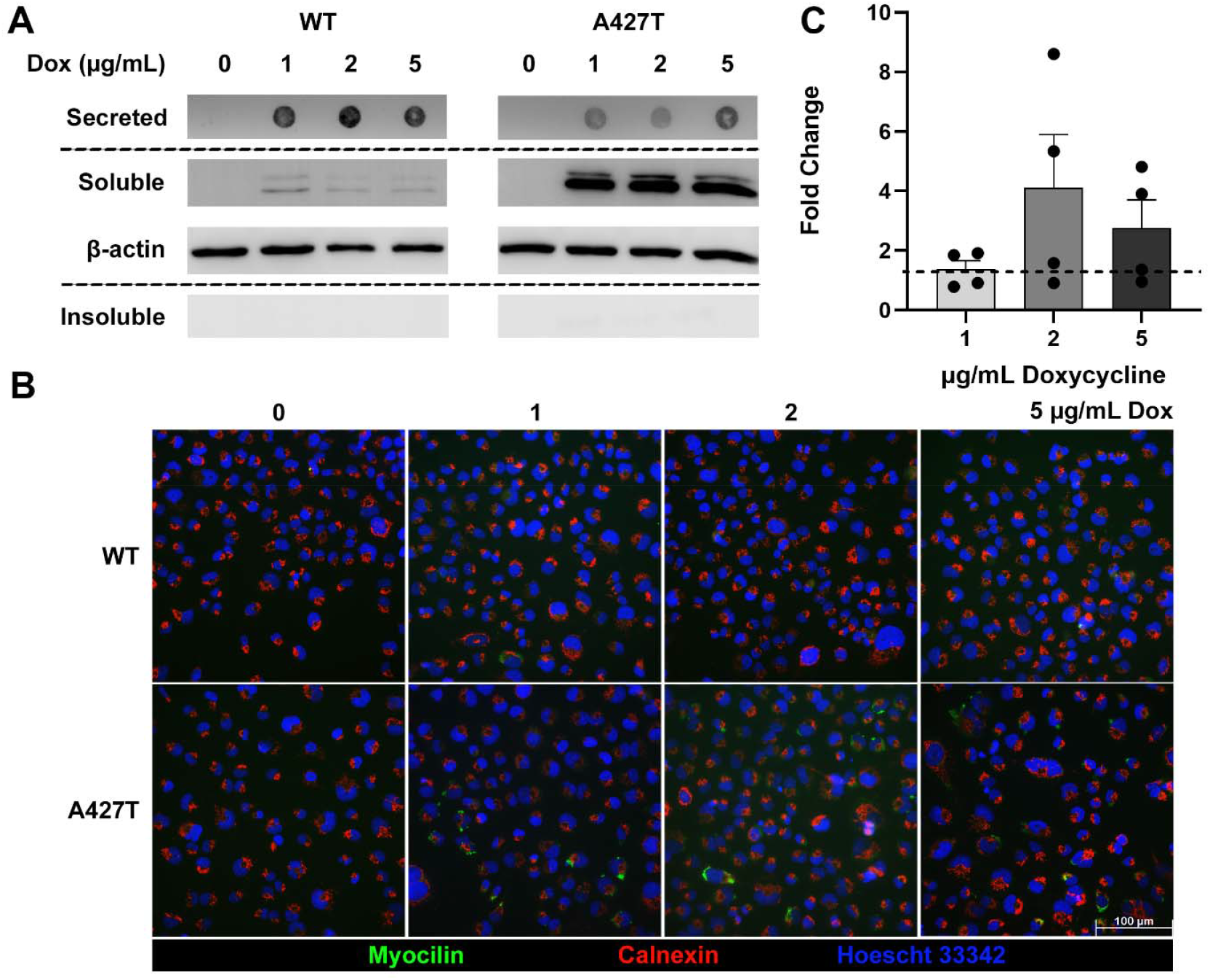
*Secretion and solubility of* myocilin^A427T^ *is intermediate between wild-type and known pathogenic mutants.* **(A)** Immunoblotting of secreted, detergent-soluble, and detergent-insoluble myocilin^WT^ and myocilin^A427T^ induced in transduced TM-1 (n=3) with increasing concentrations of doxycycline (dox). Secreted myocilin was assayed by dot blot while intracellular fractions were assessed by Western blot. β-actin serves as a load control for the detergent-soluble fraction. **(B)** Immunocytochemistry of myocilin^WT^ and myocilin^A427T^ expressed in TM-1 cells after inducing with increasing dox concentrations (n=4). Nuclei (blue); myocilin (magenta), and calnexin (green). Scale bar is 100 µm. **(C)** Quantification of (B) with fold change being myocilin ^A427T^ relative to myocilin^WT^ (dotted line). Data is mean with error bars representing s.e.m. and p > 0.05.

### Upon stress, myocilin^WT^, but not myocilin^A427T^, can be rerouted from ERAD to autophagic clearance

Prior studies of myocilin degradation confirmed involvement of the proteasomal degradation pathway (Qiu *et al*., 2014). Specifically, our lab showed involvement of Grp94 in a dominant-negative mechanism that prevented efficient retrotranslocation of the severe mutant I477N for ERAD when expressed in an inducible HEK293T cell model (Suntharalingam *et al*., 2012; Stothert *et al*., 2014; Huard *et al*., 2018). To better understand differential cellular localization and degradation of myocilin^WT^ and myocilin^A427T^, we treated transduced TM-1 cells expressing either myocilin^WT^ or myocilin^A427T^ (2 µg/mL dox, 16 hours) with proteasomal or autophagy inhibitors MG132 and bafilomycin A1 (BafA1), respectively. Cellular localization of myocilin was evaluated in immunoblot and by immunocytochemistry stained with either Calnexin to determine ER retention or atg5 to determine lysosomal recruitment **(Figure 2**).

**Figure 2.**
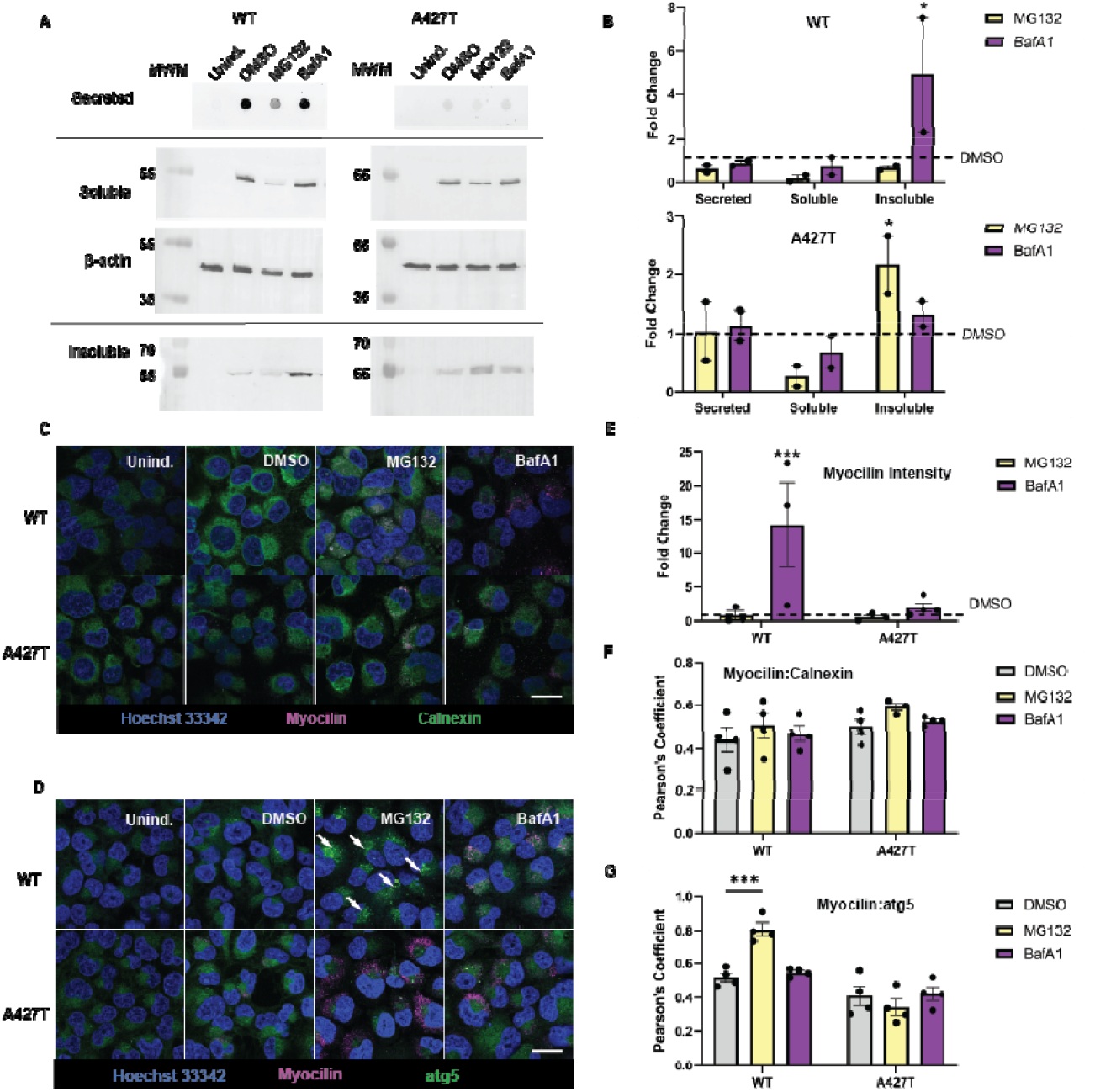
Compared to myocilin^WT^, myocilin^A427T^ cannot be trafficked efficiently to lysosomes. **(A)** Immunoblotting of secreted, detergent-soluble, and detergent-insoluble myocilin^WT^ and myocilin^A427T^ induced in transduced TM-1 and treated with MG132 and BafA1 (n=2) with uninduced controls. Secreted myocilin was assayed by dot blot while intracellular fractions were assessed by Western blot. β-actin serves as a load control for the detergent-soluble fraction. **(B)** Quantification of (A) with fold change relative to DMSO treated cells (dotted line) **(C)** Immunocytochemistry of myocilin^WT^ and myocilin^A427T^ expressed in TM-1 cells with inhibitor treatment (n=2,2). Nuclei (blue); myocilin (magenta), and calnexin (green). **(D)** Nuclei (blue); myocilin (magenta), and atg5 (green). White arrows indicate colocalization of myocilin with atg5. Scale bar is 20 µm. **(E)** Quantification of myocilin intensity relative to DMSO (dotted line) (**F**) Colocalization analysis of analysis of myocilin with calnexin in (C). (**G**) Colocalization analysis of myocilin with atg5 in (D). Error bars represent s.e.m. and *, p > 0.05; **, p>0.01; ***, p>0.001; ****, p>0.0001.

When TM-1 cells expressing modest levels of WT myocilin are stressed by MG132-induced proteasome inhibition, myocilin^WT^ does not accumulate intracellularly but rather is cleared by autophagic mechanisms. In immunoblot, compared to DMSO vehicle control, treatment with MG132 decreases secreted and intracellular soluble myocilin^WT^, but does not change the levels of detergent-insoluble (**Figure 2A, B**). Similarly, in immunocytochemistry, myocilin^WT^ is found at relatively low levels intracellularly and colocalizes with the ER marker calnexin (**Figure 2C, E, F**). Upon treatment with MG132, there is no change in calnexin colocalization but a statistically significant increase in colocalization with atg5 indicating recruitment to lysosomes (**Figure 2D, G**). Treatment with the autophagy inhibitor BafA1 shows that proteasomal clearance alone is insufficient to handle WT myocilin degradation. In Western blot, treatment with BafA1 leads to no change in secreted or soluble myocilin^WT^ but an increase in insoluble myocilin^WT^. Accordingly, in immunocytochemistry imaging there is no statistically significant change in colocalization with ER or lysosomal markers (**Figure 2C-G**). With BafA1 treatment, lysosome puncta are not observed; myocilin^WT^ is no longer localized with atg5 and myocilin^WT^ intensity increases due to inhibited autophagic degradation and subsequent intracellular sequestration (**Figure 2E**). In sum, TM-1 cells expressing myocilin^WT^ appear to handle cellular stress by engaging both proteasomal and autophagic clearance mechanisms, with the latter being the more robust degradation pathway under stress.

By contrast, in the case of myocilin^A427T^, proteasomal and autophagic mechanisms do not appear to complement one another to efficiently clear mutant myocilin. Upon treatment with proteosome inhibitor MG132, myocilin^A427T^ accumulates as a detergent-insoluble aggregate, with no change in secreted and modest reduction in soluble fractions (**Figure 2B**). Conversely, when cells were treated with autophagy inhibitor BafA1, there was no significant change in subcellular localization compared to DMSO control in Western blot (**Figure 2A**). Accordingly, in immunocytochemistry no change in colocalization was observed for either myocilin^A427T^ with calnexin (**Figure 2C and F**) nor with atg5 (**Figure 2D and G**) compared to DMSO control. These data suggest that TM-1 cells are unable to clear myocilin^A427T^ through autophagy.

In summary, we infer from these experiments that degradation of myocilin^A427T^ is primarily via ERAD and cannot be efficiently rerouted to autophagy like myocilin^WT^. Given that degradation of partially functional but reduced stability protein variants is crucial for cellular proteostasis (Flagg *et al*., 2023), it is surprising that even a mild mutation like A427T would pose a challenge for maintaining proteostasis. Still, our current findings build on the idea that promoting autophagic clearance in trabecular meshwork cells could constitute a therapeutic strategy for glaucoma.

### OLF^A427T^ structurally resembles OLF^WT^ with local perturbations near the mutation site

To better clarify the molecular origin of the aberrant cellular behavior for myocilin^A427T^ compared to myocilin^WT^, we solved the crystal structure of OLF^A427T^, to 1.45 Å resolution (**Figure 3, Table 2**). The structure confirms that the overall 5-bladed β-propeller fold of OLF, including the dinuclear metal cluster in the central hydrophilic cavity, remains intact. However, several deviations from OLF^WT^ are apparent. OLF^WT^ crystallizes with one polypeptide chain in the asymmetric subunit, in line with solution fractionation as a monomer (Huard *et al*., 2019). By contrast, OLF^A427T^ has four crystallographically independent copies of OLF^A427T^ (chains A-D, **Figure 3A**, root mean squared deviation (r.m.s.d.) = 0.207 – 0.232 Å), with loop deviations near the mutation site at residue 427 largely matching between chains A and B (blue, **Figure 3A, C, E**, r.m.s.d. = 0.232 Å) and, separately, chains C and D (pale cyan, **Figure 3A, D, E**, r.m.s.d. = 0.207 – 0.212 Å). Loop_440-446_ (green, **Figure 3B**) could not be modeled in chains A and B, whereas in chains C and D the loop could be modeled and is shifted by ∼ 5 Å. Loop_469-475_ (yellow, **Figure 3B**) could also not be modeled in chains A and B but retains WT-like configuration in chains C and D. The loss of electron density in loops of chains A and B near Thr427 is indicative of increased loop dynamics compared to OLF^WT^ but these changes do not propagate to distal parts of the protein, which was seen in other disease mutants OLF^D380A^ and OLF^I499F^ (Saccuzzo *et al*.) that elude crystallization. The collective structural perturbations likely reflect deviations noted previously in tertiary circular dichroism spectra compared to that of OLF^WT^ (Hill *et al*.).

**Table 2.** X-ray data collection and refinement statistics.

|  | **OLF<sup>A427T</sup> (PDB ID: 9DRW)** |
| --- | --- |
| **Wavelength** | 1 |
| **Resolution range** | 58.6 - 1.45 (1.50 - 1.45) |
| **Space group** | C 1 2 1 |
| **Unit cell** | 140.93 47.81 149.99 90 107.71<br>90 |
| **Total reflections** | 1084929 (73615) |
| **Unique reflections** | 160850 (12829) |
| **Multiplicity** | 6.7 (5.7) |
| **Completeness (%)** | 95.1 (76.5) |
| **Mean I/sigma(I)** | 16.2 (4.45) |
| **Wilson B-factor** | 14.7 |
| **R-merge** | 0.0641 (0.538) |
| **R-meas** | 0.0695 (0.593) |
| **R-pim** | 0.0266 (0.241) |
| **CC1/2** | 0.999 (0.912) |
| **CC\*** | 1 (0.977) |
| **Reflections used in refinement** | 160737 (12824) |
| **Reflections used for R-free** | 1999 (159) |
| **R-work** | 0.146 (0.215) |
| **R-free** | 0.188 (0.291) |
| **CC(work)** | 0.974 (0.946) |
| **CC(free)** | 0.963 (0.921) |
| **Number of non-hydrogen atoms** | 9118 |
| **macromolecules** | 8345 |
| **ligands** | 14 |
| **solvent** | 759 |
| **Protein residues** | 1029 |
| **RMS(bonds)** | 0.004 |
| **RMS(angles)** | 0.81 |
| **Ramachandran favored (%)** | 96.5 |
| **Ramachandran allowed (%)** | 3.46 |
| **Ramachandran outliers (%)** | 0.00 |
| **Rotamer outliers (%)** | 0.00 |
| **Clashscore** | 3.53 |
| **Average B-factor** | 21.2 |
| **macromolecules** | 20.0 |
| **ligands** | 32.4 |
| **solvent** | 33.6 |

Statistics for the highest-resolution shell are shown in parentheses.

**Figure 3.**
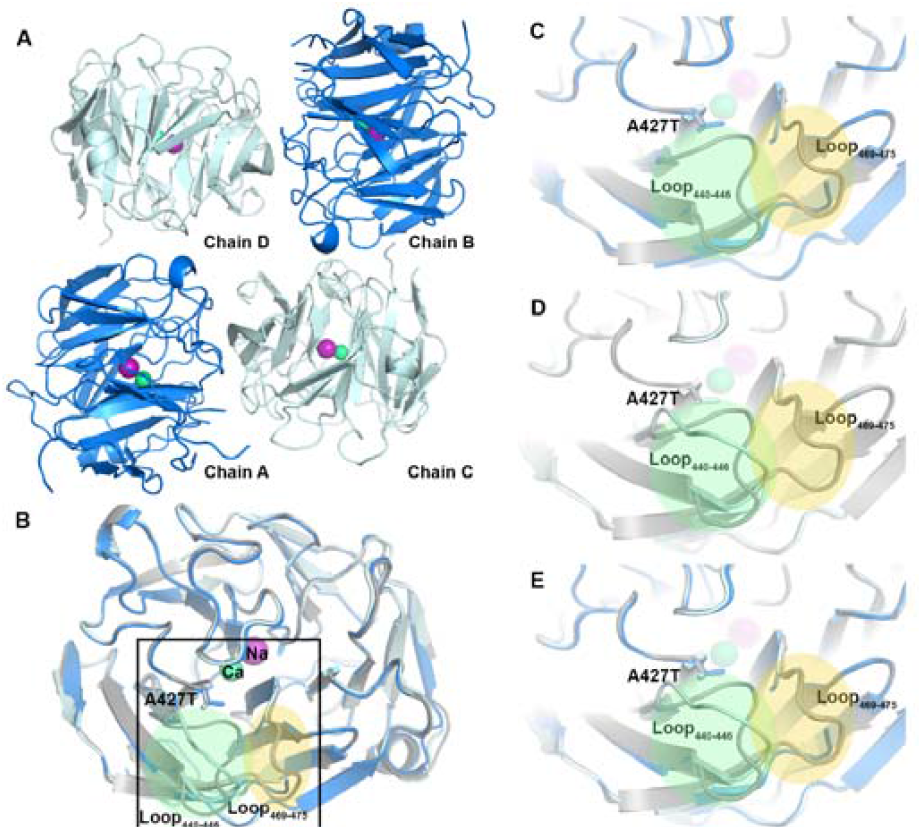
Crystal structure of OLF^A427T^. **(A)** Overall architecture of the crystal with 4 monomers in the asymmetric subunit. **(B – E)** Chains A and B (blue) have unmodeled Loop_440-446_ (green) and Loop_469-475_ (yellow) while chains C and D (pale cyan) have structured Loop_440-446_ and Loop_469-475_. The Ala427Thr mutation pushes Loop_440-446_ up and out compared to WT (PDB ID: 8FRR, gray). Loop_469-475_ is unmodeled in chains A and B while retaining a WT-like conformation in chains C and D.

### Implications for glaucoma risk

The goal to understand pathogenicity of myocilin mutations is motivated by the importance of risk assessment for glaucoma, a disease that goes unnoticed until significant and irreversible vision loss has occurred (Wang *et al*., 2019; Stein *et al*., 2021). Early monitoring for intraocular pressure elevation, which originates in the anterior eye, and subsequent intervention, are key to deterring vision loss (Stein *et al*., 2021; de Vries *et al*., 2024; Zebardast and Wiggs, 2024). However, predicting the pathogenicity of novel missense variants of myocilin is challenging.

In addition to causal disease mutations identified via Mendelian inheritance in families and clinical evaluation, myocilin variants could also contribute to the overall genetic risk without being directly causal (Allingham *et al*., 2009; Sears *et al*., 2019; Wiggs, 2022). For example, Q368X (myocilin^Q368X^), a relatively prevalent premature stop variant in myocilin (gnomAD allele frequency 1.16e-3), has been identified as a risk modifier for glaucoma (Craig *et al*., 2020; Chen *et al*., 2024). Disease causality is questioned for the myocilin^Q368X^ variant on its own due to low penetrance in population studies, but it has been shown to increase disease risk in the presence of other variants in other genes (Melki *et al*., 2003; Milla *et al*., 2013; Han *et al*., 2019; Craig *et al*., 2020; Zebardast *et al*., 2021). Myocilin^Q368X^ is currently the only myocilin variant that is included as part of a polygenic risk score developed to assess the relative risk of disease to an individual patient based on their genetic makeup (Craig *et al*., 2020).

Our findings that cellular clearance of myocilin^A427T^ is impaired despite harboring a largely WT-like biophysical and structural profile suggests that the extent of pathogenicity may be influenced by background genetics or by environmental factors. Indeed, environmental factors can make more common and low phenotypic effect variants hard to pinpoint (Manolio *et al*., 2009). For example, use of glucocorticoids can induce expression of myocilin in TM cells (Nguyen *et al*., 1998), and steroids can lead to ocular hypertension (Overby and Clark, 2015; Fini *et al*., 2017), though these effects are not thought to be directly correlated (Patel *et al*., 2017). Other environmental factors relevant to glaucoma and trabecular meshwork dysfunction that could make cells more susceptible to protein misfolding include increased reactive oxygen species and mechanical stress (Itakura *et al*., 2015; Yan *et al*., 2024). Such individualized vulnerabilities could push mild variants such as myocilin^A427T^ beyond their proteostatic threshold. In addition, currently underappreciated is the precise activation profile of UPR signaling in TM cells, which may be different from other cell types (Karagoz *et al*., 2019) and explain why mutations in myocilin are not associated with systemic disease.

There are likely other variants in myocilin that behave similarly to myocilin^A427T^ but remain obscure because genetic screening is not currently the standard of care. Therefore, our study further highlights a need for genetic screening. Furthermore, our study reiterates the critical role ERAD dysfunction and cellular proteotoxicity play in rendering myocilin mutants pathogenic, even with relatively minor structural or biophysical changes. These mechanisms should be explored further for therapeutic intervention. Overall, this study highlights the complexity of comprehending genetic risk for glaucoma.

## Materials and Methods

### Lentiviral construct for myocilin^A427T^

Site-directed mutagenesis to create the human *MYOC* A427T mutation, and insertion of wild-type and mutant cDNAs into the lentiviral vector pSLIK-hygro (Invitrogen, Carlsbad, CA) was previously described (Itakura *et al*., 2015). For packaging, lentiviral expression plasmids were co-transfected into HEK293T cells along with helper plasmids pMD.G1 and pCMVR8 (Shin *et al*., 2006). Culture medium containing viral particles was harvested after 48–72 h and stored at -70°C.

### Generation of stably transduced human trabecular meshwork cells

Immortalized TM (TM-1) cells were established from human TM cells isolated from a 30 year old donor eye as previously described using SV40 origin defective vector (Polansky *et al*., 1979; Polansky *et al*., 1984; Murnane *et al*., 1985; Filla *et al*., 2002; Faralli *et al*., 2019). Cells were validated using short tandem repeat (STR) analysis (Masters *et al*., 2001). Viral particles described above were used to infect immortalized TM-1 cells. Cell cultures were infected at a low multiplicity of infection (50 MOI) to ensure <30% infection frequency. Cells stably transduced with lentiviral expression vectors were selected by treating the cultures with 150 µg/ml hygromycin (Invitrogen, Carlsbad, CA) for 3 weeks.

For induction of myocilin^WT^ and myocilin^A427T^, TM-1 cells of stably transduced lines were seeded in 6 well plates at 200,000 cells per well. To induce the transgene, cells were then treated with dox (100–1000 ug/ml) for up to 7 days.

### Cell culture

Cells were maintained in low-glucose (1g/L) Dulbecco’s modified Eagle medium with glutamine (DMEM, Gibco) supplemented with 10% fetal bovine serum (FBS, Hyclone) and 1-2% penicillin-streptomycin (PS, Gibco) at 37 °C per consensus recommendations (Keller *et al*., 2018). Media was supplemented with 200µg/mL hygromycin (Corning) to keep cells in selection.

### Secretion Assay

Human TM-1 cells stably transduced with WT and A427T myocilin (50MOI) were seeded in 6-well plates at equivalent cell counts/replicate. The next day, the cells were treated with dox (Thermo Scientific) in serum-free media to induce myocilin expression. For dose-dependency experiments, cells were treated with 0, 1, 2, and 5 µg/mL dox for 48 hours and three separate doxycycline treatments were performed. For cellular trafficking experiments, cells were treated only with 2 µg/mL dox and two separate doxycycline treatments were performed. After 48 hours, media was removed and cells were treated with inhibitors in media at desired concentrations for 16 hours. Inhibitors included MG132 at 10µM (Sigma #M7499) and bafilomycin A1 at 100nM (Sigma SML1661-.1).

For both experiments, after treatment duration, spent media containing secreted myocilin was collected and treated with protease inhibitors cocktail containing one tablet of cOmplete EDTA-free protease inhibitor (Roche) and 1% (v/v) of phosphatase inhibitor cocktails II and III (Sigma-Aldrich). Cocktail was added at 1:100 ratio of protease inhibitor cocktail to collected media. Cells were lysed with Triton X-100 lysis buffer (100 mM Tris-HCl pH 7.4, 3 mM EGTA, 5 mM MgCl_2_, 0.5% Triton-X 100) supplemented with phenylmethylsulfonyl fluoride to final concentration of 0.6mM and protease inhibitors cocktail as described above at 1:100 ratio. Cells were lysed by rocking at 4 °C for >2 hours. Detergent-soluble (i.e., supernatant) and detergent-insoluble (i.e., pellet) fractions were separated by centrifugation. The detergent-insoluble fraction was PBS-washed before resuspending in 2X Laemmli’s Buffer + 10% 2-mercaptaethanol and sonicating with a rod sonicator (Qsonica Q125) for 5 minutes with 10-second on/off pulses at 50% amplitude. Protein concentration of the spent media and detergent-soluble fractions was determined by BCA assay (Pierce).

### Immunoblotting

In preparation for western blot analysis, detergent-soluble samples were prepared in equal protein concentration within each experiment in a final 1X Laemmli buffer containing 5-10% (v/v) β-mercaptoethanol. Samples were heated at 95°C for 5 minutes and loaded on either 12% tris-glycine gels made in house or 4–15% mini-PROTEAN TGX precast protein gels (Bio-Rad) in equal protein quantity within each fraction and experiment. Insoluble samples were heated at 95°C for an additional 15-20 minutes and spiked with additional 3 μL of β-mercaptoethanol and loaded on gels in equal volume (14 – 20 μL) per lane. For dot blot analysis, cell media were prepared in equal protein concentration within each experiment in nuclease-free water.

For Western blot analysis, gels were transferred to methanol-activated PVDF membranes using the Trans-blot Turbo transfer system (Bio-Rad). For dot blot analysis, cell media samples were deposited in equal protein quantities onto Amersham Protran nitrocellulose membranes (Sigma-Aldrich) soaked in PBS buffer (Gibco) using a Bio-Dot microfiltration apparatus (Bio-Rad). Downstream Western and dot blot steps were performed as described above. Membranes were blocked with 5% milk in PBS+0.5% Tween, washed, and incubated with 1:1000 dilution of primary antibodies (Mouse anti-myocilin, MAB3446, R&D Systems and rabbit anti-beta actin, 4970, Cell Signaling), followed by additional wash and incubation with 1:3000 dilution of secondary antibodies (anti-mouse Starbright blue 520 and anti-rabbit Starbright blue 700, Bio-Rad). After final wash blots were visualized using a ChemiDoc MP Imaging System (Bio-Rad). Results for cellular secretion assays represent at least two biological replicates.

For cellular trafficking experiments, quantification of protein bands on Western blots was performed using ImageLab (BioRad). Measurements of band density were normalized to the control sample (no induction) within each experimental treatment. Statistical significance was determined with one-way ANOVA with GraphPad.

### Immunocytochemistry

For dose-dependency experiments, human TM-1 cells stably transduced with myocilin^WT^ and myocilin^A427T^ myocilin (50MOI) were seeded on glass coverslips (12 mm diameter, Fisher) coated with 2% gelatin (Sigma-Aldrich) in 24 well plates. The next day, cells were treated with 0, 1, 2, and 5µg/mL doxycycline (n=4) to induce myocilin expression. Three separate replicate inductions were performed. After 48 hours, cells were washed with 1X PBS (Gibco), fixed with 10% formalin (Fisher Healthcare), permeabilized with 0.03% Triton X-100 (VWR Health Sciences), and blocked in 5% milk before incubating with primary antibodies >12 hours (1:1000 rabbit anti-myocilin, Abcam Ab41552; 1:1000 mouse anti-calnexin, Invitrogen MA3-027). Following a PBS wash, cells were incubated with 1:1000 secondary antibodies (goat anti-rabbit IgG, Alexa Fluor 488, Invitrogen A11034; goat anti-mouse IgG, Cyanine5, Invitrogen A10524) for 1 hour. Following another PBS wash, cells were incubated with 1:1000 Hoechst 33342 (catalog number ENZ51035K100, Enzo Life Sciences, Farmingdale, NY, USA) for ∼30 minutes. Cells were washed before fixing coverslips with ProLongTM Diamond Antifade Mountant (Invitrogen) and allowed to cure for at least 24 hours. Negative controls for each condition were treated with only the secondary antibodies and not the primary antibodies to eliminate non-specific antibody detection. Two to three positively and 1-3 negatively stained coverslips were prepared for each condition and 3-4 representative images were acquired per coverslip at 40X magnification using a Leica DMB6 fluorescent microscope (Leica Microsystems). Brightness and contrast of all images in the panel was equivalently adjusted with Leica Application Suite X software and Adobe Photoshop. Quantification was conducted using ImageJ (Schneider *et al*.) by measuring the mean gray value for both MYOC and Hoescht 33342 fluorescence after autothreshholding with the Triangle autothreshholding method. MYOC fluorescence values were normalized to Hoescht 33342 fluorescence values before averaging per replicate and condition. Average values were background-subtracted with the average value for 0 µg/mL control for myocilin^WT^ or myocilin^A427T^. From these background-subtracted, normalized values, fold change was calculated for each A427T treatment concentration relative to WT. Outlier data was excluded by the interquartile method. Data were analyzed by multiple unpaired, two-tailed t-tests (myocilin^WT^ versus myocilin^A427T^ within corresponding treatments) using GraphPad Prism.

For cellular trafficking experiments, human TM-1 cells stably transduced myocilin^WT^ and myocilin^A427T^ (50MOI) were seeded on glass coverslips (12 mm diameter, Fisher) coated with 1.5 µg/mL PLL (ScienCell Catalog No. 0403) in 24 well plates. The next day, the cells were treated with 2 µg/mL doxycycline (n=2) independent experiments. After 16-hour drug treatment cells were washed with 1X PBS (Gibco), fixed with 10% formalin (Fisher Healthcare), permeabilized with permeabilization buffer containing 0.05% Triton X-100 (VWR Health Sciences), and blocked in 2% BSA overnight before incubating with primary antibodies for 4 hours (1:200 mouse anti-myocilin, R&D Myocilin MAB3446; 1:200 rabbit anti-calnexin, ThermoFisher PA5-34754). Cells were incubated with 1:1000 secondary antibodies (goat anti-rabbit IgG, Alexa Fluor 488, Invitrogen A11034; goat anti-mouse IgG, Cyanine5, Invitrogen A10524) for 1 hour. Following another PBS wash, cells were incubated with 1:1000 Hoechst 33342 (catalog number ENZ51035K100, Enzo Life Sciences, Farmingdale, NY, USA) for ∼30 minutes. Cells were washed before fixing coverslips with ProLongTM Diamond Antifade Mountant (Invitrogen) and allowed to cure for at least 24 hours. Coverslips were prepared for each condition including uninduced cells of two biological replicates containing two technical replicates (coverslips) with 3-4 representative images were acquired per coverslip. Images collected at 63X with Zeiss 700 Laser Scanning Microscope with same laser power conditions. Image channel brightness and gamma were equivalently adjusted with the Zen Blue software. Intensity quantification was conducted measuring the mean gray value of adjusted images and colocalization analysis was conducted using the JACoP plugin (Bolte and CordeliÈRes, 2006) in ImageJ (Schneider *et al*.). Statistical analysis was performed by one-way ANOVA using GraphPad Prism.

### OLF expression and purification

OLF^A427T^ was recombinantly expressed and purified as described previously (Donegan *et al*., 2015) minor modifications. Briefly, Rosetta-gami 2(DE3) cells (Novagen) transformed with plasmids were grown in Superior broth (US Biological) supplemented with 60 µg/ml of ampicillin and 34 µg/ml of chloramphenicol. Cultures were induced at an OD600 of 1.5 at 18°C with 0.5 mM isopropyl-β-D-thiogalactopyranoside and 100 mM CaCl_2_, and cells were allowed to grow for 16 h. The cells were pelleted, flash frozen in liquid nitrogen, and stored at −80°C. Protein was purified by amylose affinity chromatography and size-exclusion chromatography as reported previously (Donegan *et al*., 2012). Purified OLF^A427T^ was stored in PGF buffer (10 mM Na_2_HPO_4_/KH_2_PO_4_, 0.2 M NaCl, pH 6.8). Purity was assessed by standard 12% SDS-PAGE analysis using stain-free gels visualized with the Bio-Rad ChemiDoc MP imaging System.

### Crystallization and structure determination

Purified OLF^A427T^ (in 10 mM HEPES, pH 7.2 with 200 mM NaCl) was concentrated to 10 mg/ml. Crystals were grown by the hanging-drop method by equilibration against a reservoir containing 10% PEG 8000 and 0.05 M MgCl_2_. Crystals were cryo-cooled in a solution containing reservoir solution supplemented with 30% glycerol. X-ray diffraction datasets were collected on the NYX beamline 19-D at Brookhaven National Laboratory National Synchrotron Light Source II (NSLS-II) and were processed using HKL-2000 (Otwinowski and Minor, 1997). Structures were solved by molecular replacement using Phaser (McCoy, 2007), using OLF^WT^ (PDB ID: <u>4WXQ</u>) (Donegan *et al*.) as the search model. The models were iteratively refined using Coot (Emsley *et al*., 2010) and Phenix.refine (Afonine *et al*., 2012). Figures were prepared in PyMOL. The structure was deposited in the PDB with accession code 9DRW. Data collection statistics are reported in **Table 2**.

## Abbreviations

myocilin^A427T^: Myocilin A427T mutant
ER: endoplasmic reticulum
ERAD: ER-associated degradation
UPR: unfolded protein response
IOP: intraocular pressure
TM: trabecular meshwork
POAG: primary open angle glaucoma
(JOAG): juvenile open angle glaucoma
OLF: olfactomedin
OLF^A427T^: A427T OLF mutant
OLF^WT^: wild-type OLF
T (ThT): thioflavin
dox: doxycycline
myocilin^WT^: wild type myocilin
A1: bafilomycin BafA1
r.m.s.d.: root mean squared deviation
(PBS): phosphate buffered saline
OD: optical density
PGF buffer: 10 mM Na_2_HPO_4_/KH_2_PO_4_, 0.2 M NaCl, pH 6.8
WT: wild-type

## Acknowledgements

The authors acknowledge use of the core facilities at the Petit Institute of Biosciences and Bioengineering. This work was funded by NIH R01EY021205 (to RLL). HAY was funded in part by T32EY007092 and F32EY03628. HFS was supported in part by 5T32EY007092. KRH was supported in part by US Department of Education GAANN grant P200A210014. Access to NYX beamline 19-D at the National Synchrotron Light Source II (NSLS-II) was provided through the Southeast Regional Collaborative Access Team (SER-CAT). SER-CAT is supported by its <u>member institutions</u>, equipment grants (S10_RR25528, S10_RR028976 and S10_OD027000) from the National Institutes of Health, and funding from the Georgia Research Alliance. NSLS-II is a U.S. Department of Energy (DOE) Office of Science User Facility operated for the DOE office of Science by Brookhaven National Laboratory under contract No. DE-SC0012704. Use of the NYX beamline 19-ID at the was supported by the New York Structural Biology Center. NYX detector instrumentation was supported by grant S10OD030394 through the office of the Director, National Institutes of Health.

## Notes

### Competing Interest Statement

The authors have declared no competing interest.

